# Differential tolerance of *Zymoseptoria tritici* to altered optimal moisture conditions during the early stages of wheat infection

**DOI:** 10.1101/867572

**Authors:** Anne-Lise Boixel, Sandrine Gélisse, Thierry C. Marcel, Frédéric Suffert

## Abstract

Foliar plant pathogens require liquid or vapour water for at least part of their development, but their response and their adaptive tolerance to moisture conditions have been much less studied than other meteorological factors to date. We examined the impact on the wheat-*Zymoseptoria tritici* interaction of altering optimal moisture conditions conducive to infection. We assessed the responses *in planta* of 48 *Z. tritici* strains collected in two climatologically distinct locations (Ireland and Israel) to four high moisture regimes differing in the timing and the duration of uninterrupted exposure to saturated relative humidity (100% RH) during the first three days of infection. Individual- and population-level moisture reaction norms expressing how the sporulating area of a lesion change with the RH conditions were established based on visual assessments of lesion development at 14, 17 and 20 days post-inoculation (dpi). Our findings highlighted: (i) a critical time-dependent effect on lesion development of uninterrupted periods of exposure to 100% RH during these earliest infection stages; (ii) a marked interindividual variation in the sensitivity to RH conditions both in terms of strain average moisture response and plasticity; (iii) a higher tolerance – expressed at 14 dpi, not later – of the Israeli population to early interruption of optimal moisture conditions. By indicating that sensitivity to sub-optimal moisture conditions may vary greatly between *Z. tritici* individuals and populations, this study highlights the evidence of moisture adaptation signature in a plant pathogen. This suggests that understanding such variation will be critical to predict their response to changing climatic conditions.

## Introduction

Micrometeorological factors within the plant canopy define the physical environment in which foliar crop pathogens develop (Campbell & Norman, 1998). Temperature and moisture are key factors determining the outcome of host-pathogen interactions with critical effects highlighted at each epidemiological stage of foliar diseases. The presence, abundance and persistence of moisture determine whether the biological processes of infection, and colonization, accelerate, slow or stop altogether, with consequences for the incubation and latent periods, spore production and release (Huber & Gillespie, 1992). The effects of temperature have been quantified in detail, but much less is known about the precise impacts of moisture levels and thresholds, whether considered in terms of relative humidity (RH) and leaf wetness (presence of water – films and droplets – on leaf surfaces, caused by precipitation, irrigation, guttation or dew), on pathogen and disease development. Liquid water appears on leaf surfaces when its temperature is less than the dew point temperature of the air (Weiss, 1990). The impact of RH vs. leaf wetness has been little addressed by plant pathologists likely because of both conceptual and technical difficulties. This gap in knowledge and practice may reflect the experimentally greater challenge of controlling, measuring and reporting moisture conditions with sufficient resolution and fidelity (Rowlandson *et al.*, 2015). Indeed, moisture conditions display a high degree of spatio-temporal variability in the field as they result from the dynamic equilibrium between water interception and evaporation. The accurate control of moisture conditions requires controlled environment plant growth chambers in which fluctuations in other meteorological variables are minimized. For instance, RH is temperature-dependent as it is defined as the ratio of the partial pressure of water vapor to the equilibrium vapor pressure of water at a given temperature. Leaf wetness duration (LWD), by contrast, is even less easy to define (no single meteorological definition; Dawson & Goldsmith, 2018), lacks of a standard for its measurement (Sentelhas *et al.*, 2004) and exhibits heterogeneous spatio-temporal patterns as different parts of the leaves may be wet and dry at different times (Huber & Gillespie, 1992). “Do pathogens respond to the absolute quantity of water vapour held in the atmosphere or to a relative percentage?” is a question that is difficult to answer. Indeed, as RH is temperature-dependent, it cannot be theoretically compared unless temperature regimes (and so water carrying capacity) are exactly the same.

Several epidemiological studies have identified prolonged wetness as an important factor for the success of infection, and several disease-forecasting models contain the variable LWD (Rowlandson *et al.*, 2015). Although fewer studies have focused on RH requirements for optimal infection, some have established that the RH threshold (the level below which infection does not occur) differs between types of pathogens, up to very high RH requirements for some fungal species (e.g. 90% for *Magnaporthe oryzae*, Li *et al.*, 2014; 95% for *Didymella rabiei*, Jhorar *et al.*, 1998).

Studies of the response of a fungal pathogen to moisture conditions at the leaf scale during the earliest stages of infection (incubation period) are particularly relevant, because this period, together with the spore release and dispersal stages, is one of the most sensitive to LWD and RH in the vast majority of pathogens. The causal agent of Septoria tritici blotch, *Zymoseptoria tritici*, is a highly relevant biological model in this context because of: (i) its long latent period, spanning from 10 to 20 dpi (days post-inoculation) on seedlings and 15 to 30 on adult plants under controlled conditions at an average daily temperature of 18°C (Suffert & Thomson, 2018) *vs*. 15 to 35 dpi in field conditions, depending on temperature (Shaw, 1990); (ii) the crucial impact of moisture – RH and leaf wetness – at all stages of the disease cycle in field conditions (Shaw, 1990).

The impacts of the duration and interruption of the wet period on *Z. tritici* (post-inoculation LWD) have already been investigated in controlled conditions (Holmes, 1974; Shearer & Zadoks, 1972; Eyal, 1977; Hess *et al.*, 1987; Chungu *et al.*, 2001; Magboul *et al.*, 1992; Shaw, 1990; Shaw, 1991; Fones *et al.*, 2017). An increase in RH favors an increase in overall infection rate (penetration of leaf tissues via the stomata), hyphal growth and pycnidiation intensity (Shaw, 1991; Fones *et al.*, 2017). RH has been inferred to be optimal between 3 and 4 days post-inoculation, at 100% (Chungu *et al.*, 2001; Suffert *et al.*, 2013), and pycnidiation has been reported to occur at a RH of 35-100% (Pachinburavan, 1981). However, threshold RH levels have not been experimentally tested, except for 50%-75%-100% comparisons (Shaw, 1991). The RH requirement threshold for *Z. tritici* development therefore remains unknown. LWD has also been shown to have critical effects on infection, pycnidiation and, in some cases, the duration of the latent period (Chungu *et al.*, 2001; Shaw, 1990). These experimental results are supported by the importance of the presence and duration of moisture (particularly LWD) for the prediction of Septoria tritici blotch epidemics in wheat (Hess & Shaner, 1987).

Reviews on the impact of climate change on plant diseases have stressed the importance of focusing on changes in temperature and rainfall patterns (Garrett *et al.*, 2006). Global climate models (GCMs) predict more frequent and extreme rainfall events and higher atmospheric water vapor concentrations with increasing temperature (Huntingford *et al.*, 2003). Until recently, it was difficult to obtain LWD and RH, which critically influence plant pathogen infection and disease development, from GCM outputs (Chakraborty *et al.*, 2000). It had been suggested that higher levels of canopy moisture promoted the development of a range of foliar pathogens, but this factor was never considered essential and was rarely taken into account in discussions on the effects of climate change on plant diseases (e.g. West *et al.*, 2012). However, the changes in RH are likely to be at least as marked as those in temperature in semi-arid zones, such as the area around the Mediterranean basin. The predicted slight decrease in the duration of wet periods and slightly warmer conditions will probably have opposite effects, decreasing and promoting infection, respectively, and detailed modeling approaches are therefore required to predict the overall outcome. This paradigm shift will require an understanding of the diversity of responses and levels of adaptation of pathogen populations to contrasting RH conditions. This understanding can be developed by identifying differences in moisture responses between pathogen genotypes within the same species, and, more generally, by characterizing the intra-*vs.* inter-population phenotypic variability connected to genetic differentiation at neutral marker loci to reveal local adaptation (Merilä & Crnokrak, 2001).

RH may have been much less considered than temperature to date because there are more convenient methods for thermal phenotyping (e.g. Bernard *et al.*, 2013; Boixel *et al.*, 2019a) than for moisture phenotyping (e.g. Li *et al.*, 2014; Xu *et al.*, 2016). Assessments of the differences in response to temperature *in vitro* (tests conducted without interaction with the plant) are biologically meaningful but have to be compared with *in planta* responses (tests conducted in interaction with the plant), as done for instance by Boixel *et al.* (2019a). By contrast, differences in the response to humidity must be performed *in planta*, particularly for the early stages of infection.

In this study, we investigated the diversity in moisture responses in *Z. tritici*, at the individual and population levels, for two field populations collected from contrasting climatic zones (humid *vs*. dry environment). We evaluated the aggressiveness of the individuals of these populations on wheat seedlings subjected to four moisture regimes. These regimes were defined by increasingly altering the inferred optimal RH conditions for *Z. tritici* over the first three days after inoculation that were reported as critical in the literature. This was done by sequentially incorporating interrupted periods of exposure to 100% RH resulting in the study of diurnal changes in humidity (reflecting what is happening in the field) rather than different levels of constant humidity.

## Methods

### Fungal material

Two *Z. tritici* populations were collected on wheat after the stem elongation stage from contrasting Köppen-Geiger Euro-Mediterranean climatic zones: (i) an Irish population (IR) sampled in July 2016 on cultivar JB Diego from a single field at Carlow (52°83’65’’ N - 6°93’41’’ E; Cfb, oceanic climate); (ii) an Israeli population (IS) sampled in March 2017 on cultivar Galil from a single field at Kiryat-Tivon (32°71’62’’ N - 35°12’74’’ E; Csa, hot-summer Mediterranean climate). These two populations, which have already been characterized for thermal adaptation and genotyped for 12 neutral genetic markers (Boixel *et al.*, 2019b), originate from environments with different average air moisture conditions, mainly due to marked differences in annual temperature and rainfall regimes (Figure 1). We randomly selected 24 IR and 24 IS *Z. tritici* isolates from these populations and checked that they were different multilocus genotypes (Boixel *et al*., 2019b). These populations were phenotyped in three sequential randomized batches of 16 isolates at a time under controlled conditions in a growth chamber experiment with four treatments (different moisture regimes but identical temperature conditions). For each batch, each treatment (isolate × RH conditions) was represented by six replicate wheat seedlings. These six replications were split in two groups of three each that were randomly assigned at opposite ends of the growth chamber to avoid confounding of treatment and growth chamber effects (potential impact of uncontrolled spatial variations in microclimatic conditions in the chamber; Potvin *et al.*, 1990). Thus, we used a randomized complete block design with locations in the growth chamber as blocks, isolate and RH conditions as treatments, and plants as replications.

**Fig. 1.**
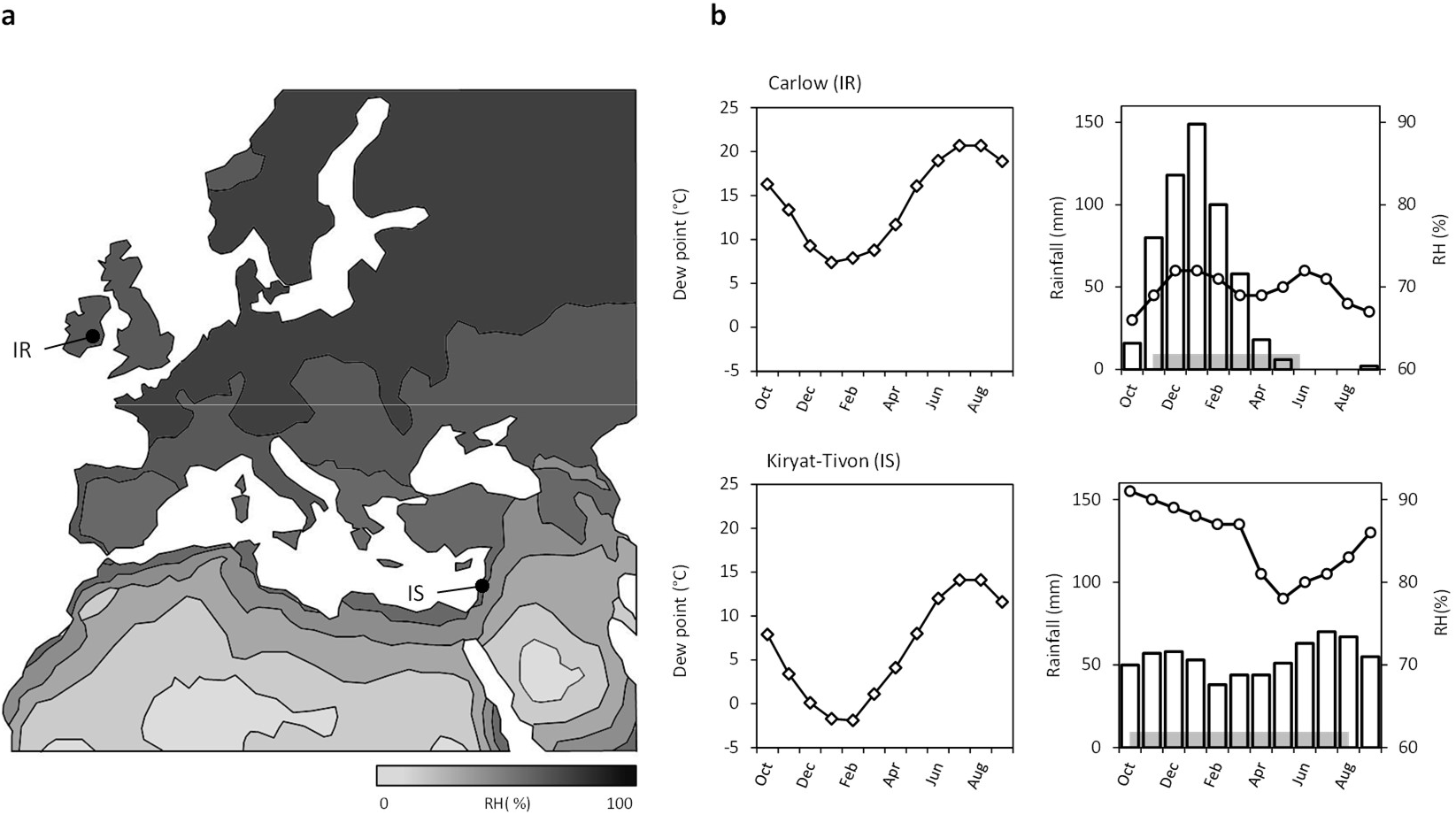
Relative humidity (RH) conditions and precipitation at the two sampling sites. (a) Average annual air RH in the Euro-Mediterranean area (data from the Atlas of the Biosphere, http://atlas.sage.wisc.edu/); (b) average monthly air RH (circles), rainfall (bars) distribution (1982-2012 data from the World Climate Database, https://en.climate-data.org/), and dew point (diamonds) calculated using monthly air RH and temperature values (https://www.calculator.net/dew-point-calculator.html) at the sites from which the two *Zymoseptoria tritici* populations of 24 isolates each were collected (Carlow in Ireland for the IR population; Kyriat-Tivon in Israel for the IS population). The horizontal gray bar indicates the standard wheat-growing period (from sowing to harvest).

### Plant material

The 48 isolates were phenotyped on the highly susceptible bread wheat cultivar ‘Taichung 29’, commonly used as a susceptible check cultivar in the screening of wheat germplasm and in tests of global panels of *Z. tritici* isolates (e.g. Makhdoomi *et al.*, 2015). Wheat seeds were sown in 0.4-liter pots, so as to obtain three seedlings per pot. The pots were kept in a growth chamber, under a controlled light/dark cycle (16 hours of light at 300 μmol.m^−2^.s^−1^ - Osram Lumilux L58W/830 - at 22°C and 80 % RH / 8 hours of dark at 18 °C and 100 % RH). These conditions were kept the same throughout the duration of the experiment.

### Inoculation procedure

Isolates were grown in Petri dishes containing PDA (potato dextrose agar, 39 g L^−1^) at 18°C, in the dark, for five days. Spores suspensions were prepared by flooding the surface of the five-day-old cultures with sterile distilled water and then scraping the agar surface with a sterilized glass rod to release the yeast-like spores (also called ‘conidia’ or ‘blastospores’). The concentration of spores in the suspension was adjusted to 10^5^ spores.mL^−1^, which is more relevant here than much higher concentrations because it maximises differences in disease expression and facilitates the detection of the effect of an experimental factor. We added one drop of Tween 20 (Sigma) per 15 mL suspension (0.1 % v/v). Each spore suspension was inoculated with a paintbrush over a 7.5 cm mid-length portion of the first true leaves of six 16-day-old seedling plants.

### Differential exposure to moisture regimes during the incubation period

After inoculation, the six plants from each isolate × RH condition interaction were split into two identical groups and assigned to two opposite ends of the growth chamber and thus considered as two blocks. Plants were subjected to four different moisture regimes (R0, R1, R2, and R3) defined based on the alteration of the reported optimal conditions for infection over the first three days after inoculation (the time required for pycnidiospore germination, epiphytic hyphal growth and stomatal penetration; Duncan & Howard, 2000). Indeed, most *Z. tritici* inoculation protocols include a bagging time of three days at 100 % RH (Chungu *et al.*, 2001; Suffert *et al.*, 2013). As such, moisture conditions were modulated by varying the duration of time after inoculation (0, 1, 2 and 3 days, respectively) at which inoculated plants were continuously subjected to 100% RH. Saturated RH was reached when plants were covered with transparent polyethylene bags previously moistened with distilled water. Uncovered plants were exposed to the diurnal RH changes of the growth chamber (RH of 100% in the dark and of 80% during light periods; average automatic recording, every 15 minutes, with a ventilated sensor). Water condensation was maintained on the bags during the night at 18°C and the day at 22°C but liquid water was not observed on the leaves, while theoretically possible (100% RH at 18°C gives a dew point of 18°C and 100% RH at 22°C gives a dew point of 22°C), maybe due to a slight difference between temperatures of leaf and bag surface. This is likely because the leaves were not radiatively cooled by exchange with sky as they would be in field conditions (especially on clear nights) but rather minutely warmed by exchanges with the walls of the growth chamber. Liquid water was not observed on the uncovered leaves during the night while some water condensation appeared on the walls (100% RH at 18°C gives a dew point of 18°C) and the dew point was out of reach during the day (80% RH at 22°C gives a dew point of 18.4°C).

Concretely, moisture regimes consisted in four durations of time at which infected plants were bagged (Figure 2): no-bagging treatment (R0); one-day-bagging treatment (R1); two-day-bagging treatment (R2); three-day-bagging treatment (R3). To establish moisture reaction norms for characterization and comparison of the RH sensitivity of the 48 isolates (see ‘*Data analyses’* section below), we summarized the lowest to highest moisture regimes in terms of the mean RH prevailing during the three-day-post-inoculation period for ease of calculation: 88.3% (R0), 92.2% (R1), 96.1% (R2) and 100% (R3). We chose not to characterize each moisture regime by the duration of time for which RH = 100 %. Indeed, there is no evidence that a RH of 80 %, which is higher than the RH value below which infection is thought to be impossible (RH = 50%; Shaw, 1991) but below the lowest humid conditions tested here, can prevent fungal growth and stop infection.

**Fig. 2.**
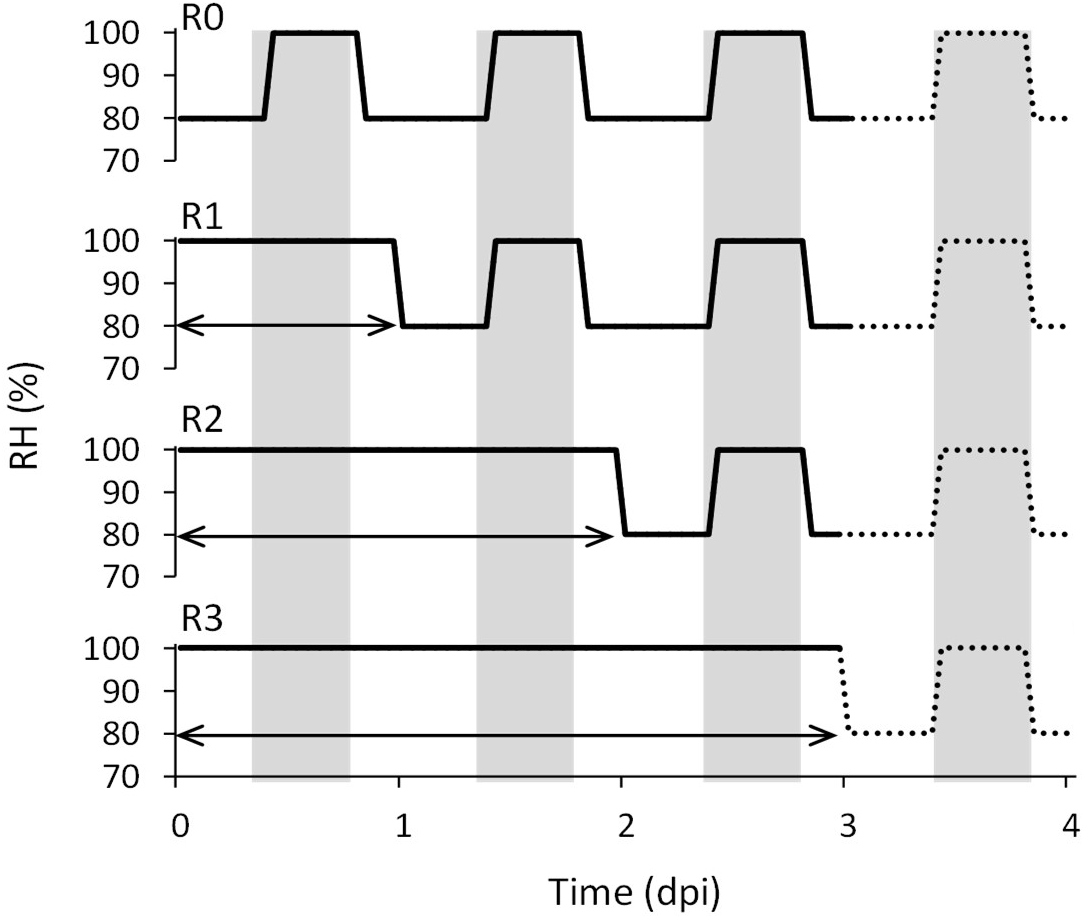
Moisture regimes exerted over the three first days post-inoculation (dpi). Solid (from 0 to 3 dpi) and dotted (after 3 dpi) lines depict the four post-inoculation moisture regimes to which the wheat seedlings were subjected in the early stages of infection: R0 = no-bagging treatment; R1 = one-day-bagging treatment; R2 = two-day-bagging treatment; R3 = three-day-bagging treatment. White and gray areas indicate successive light and dark periods, respectively. Horizontal arrows indicate the period during which wheat plants were enclosed within polyethylene bags to maintain 100% RH.

### Disease assessment

Disease severity was assessed by eye, by the same assessor, at 14, 17 and 20 dpi, as the percentage of the inoculated leaf surface (0, 1, 2, 3 and 5%, then increments of 5% up to 100%) displaying visible fruiting bodies or pycnidia (sporulating area). The robustness of this visual assessment method was established in previous studies performed on both seedling and adult wheat plants under similar facilities (Suffert *et al.*, 2013; 2016), as widely used in phytopathometry.

### Data analysis

#### Effect of moisture conditions on disease severity

The experimental and biological main effects on the variability of sporulating area (SPO, expressed as a percentage of the inoculated leaf surface) and their interactions were assessed by a statistical modeling approach. The variance of sporulating area measured on six replicates per treatment per isolate was divided into sources attributable to: series corresponding to the three randomised batches of 16 strains each (S), block (B), RH conditions (H), population (P), isolate I(P), RH conditions × population interaction (H×P), RH conditions × isolate interaction (H×I(P)), residuals (ε_sbpih_). We analyzed these deviance components, by fitting a generalized linear model (GLM) with a log-link function and quasi-Poisson errors to account for overdispersion, according to the following model (R *glm* function): SPO_sbpih_ = S + B + P + I(P) + H + H×P + H×I(P) + ε_sbpih_. Tukey’s HSD *post-hoc* comparisons of differences in SPO means (multivariate analysis of variance) and homoscedasticity (Levene’s test) between IR and IS populations were performed to investigate differences in inter- and intra-population responses to moisture conditions.

#### Establishment of individual moisture reaction norms

Individual moisture reaction norms, describing the pattern of SPO as a function of relative humidity, were established for each isolate. Reaction norms were assumed to be linear within the investigated 80-100% RH range and were therefore estimated by a linear regression SPO = a × RH + b. Individual reaction norm properties were summarized by three parameters accounting for sensitivity to RH: (i) the intercept ‘*b’* corresponding to the elevation or the sensitivity to the driest environment; (ii) the steepness of the slope ‘*a’* corresponding to the response to variation, *i.e.* whether phenotypic responses increased or decreased in extreme environments and to what extent; (iii) the midpoint ‘y_94.2%_’ corresponding to the average response or the response at the midpoint of the RH range, to consider shifts in the entire reaction norm. The heterogeneity of RH sensitivity across isolates at the individual level and the difference between IR and IS population-level moisture reaction norms were assessed on the basis of these three parameters. We did not account for the variation in the goodness-of-fit of individual reaction norms for the population-level analysis, as routinely employed when carrying a linear reaction norm approach (de Jong, 1995).

#### Differentiation in individual and population responses

Phenotypic differentiation in terms of the intercept, slope and midpoint was compared to between-population neutral genetic differentiation, to determine the potential for local adaptation of moisture responses (P_ST_ - F_ST_ comparisons performed with the ‘*Pstat*’ R package). Neutral genetic variability, and the structure and distribution of diversity between and within the IR and IS populations were characterized in a previous study (Boixel *et al.*, 2019b) based on microsatellite genotyping data acquired for 12 SSRs (ST1, ST2, ST3A, ST3B, ST3C, ST4, ST5, ST6, ST7, ST9, ST10, ST12 neutral microsatellites; Gautier *et al.*, 2014). In parallel, components of phenotypic variation in the response to the four moisture regimes (R0, R1, R2, R3) were extracted by principal component analysis. We projected all four-fold phenotypes onto a two-dimensional space and clustered the isolates (hierarchical classification on principal components, HCPC) on the basis of their sensitivities to the four moisture regimes (‘*FactoMineR*’ and ‘*factoextra*’ R packages) to offer an integrated view of the responses to the four RH conditions (R0, R1, R2, R3) and to better distinguish singular phenotypes.

## Results

### Overall time-dependence effect of the experimental conditions

The series (S) and block (B) effects were significant despite the strong control over RH conditions of our experimental design (saturation-level application), consistently with the spatial and temporal variability of other micrometeorological conditions reported to occur within growth chambers, even under controlled conditions (e.g. gradient in irradiance intensity due to artificial lighting; Poorter *et al.*, 2012). Mean RH (H) had a significant effect on sporulating area at all times of disease assessment (14, 17 or 20 dpi; GLM, *p*-value < 0.01; Table 1, Figure 3). The significant differences in sporulating area induced by different *Z. tritici* isolates (I(P)) confirmed the genetic origin of the variability in aggressiveness. The population effect (P) was significant at 14 dpi, when sporulating area was greater for IS than for IR isolates under the R0, R1 and R2 regimes (Figure 4). Significant H×I(P) and H×P interactions highlighted differential individual and population-level tolerance to the four moisture regimes (Table 1). At 14 dpi, differences in the mean sporulating area (GLM *post-hoc* analysis of the H×P interaction, *i.e.* the interaction of moisture regimes and differential population-level response to these conditions) and its intrapopulation variance (Levene’s test for homogeneity of variance) between the IR and IS populations were significant (*p*-value < 0.05; Figure 4). These differences diminished at 17 dpi (not statistically significant) and completely disappeared by 20 dpi.

**Table 1.**
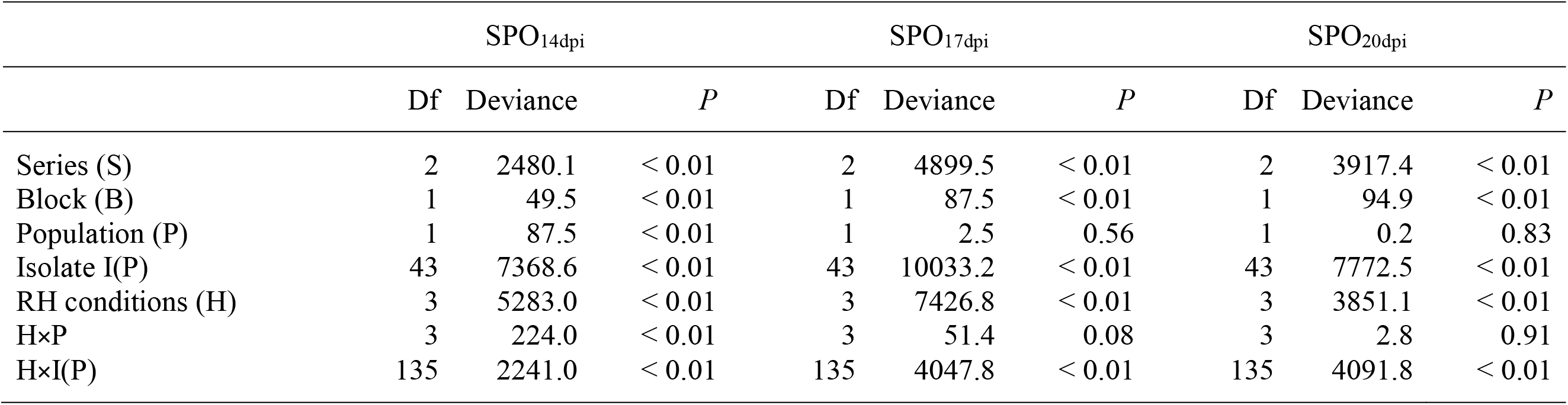
Analysis of the deviance of sporulating area (SPO in %) induced by 48 *Zymoseptoria tritici* isolates on wheat seedlings exposed to four high moisture regimes (Figure 2) at 14, 17 and 20 dpi. A quasi-Poisson generalized linear model (GLM) with a log-link function was fitted to experimental data, to assess the relative importance of the factors considered for the observed variation: series (S), block (B), RH conditions (H), population (P), isolate I(P), RH conditions × population interaction (H×P), RH conditions × isolate interaction (H×I(P)). The chi-squared test statistics reported in the table correspond to the change in degrees of freedom (Df), the components of deviance (Deviance) and their corresponding *p*-values (*P*).

**Fig. 3.**
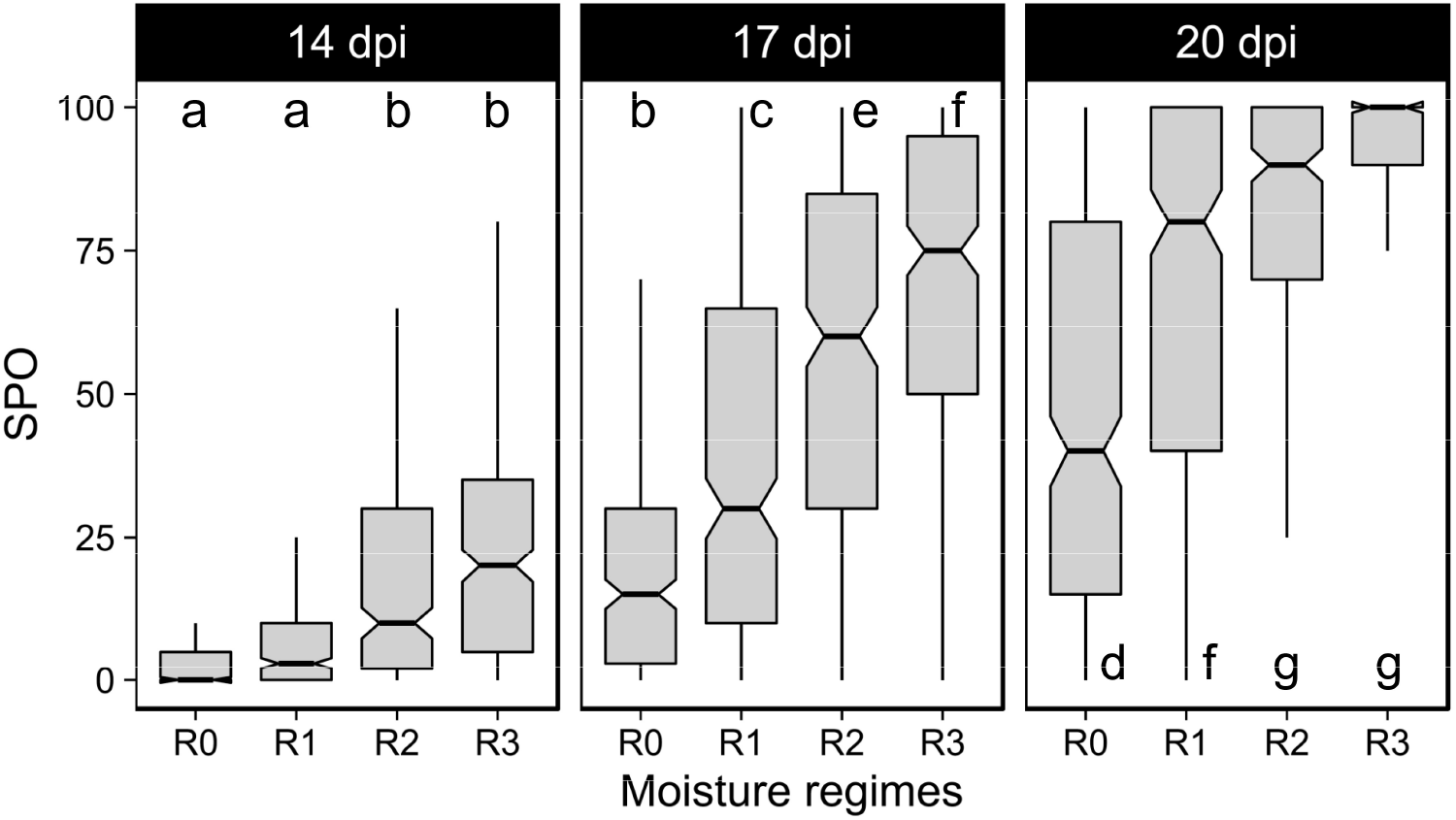
Effect of moisture regime on the percentage of sporulating area (SPO) induced by *Zymoseptoria tritici*, by disease assessment time (14, 17, 20 dpi). Data are means of 48 isolates (*n* = 48) where each isolate is described by SPO assessments conducted on 6 wheat leaves taken from 2 pots placed in different growth chamber positions (opposite ends). Medians are indicated by the lines at which the notches converge. Different letters indicate significant differences in SPO between experimental conditions (combination of moisture regimes and disease assessment time) in pairwise comparisons using Tukey HSD *post-hoc* test comparisons (*p*-value < 0.05; multivariate analysis of variance). Moisture regimes: R0 = no-bagging treatment; R1 = one-day-bagging treatment; R2 = two-day-bagging treatment; R3 = three-day-bagging treatment.

**Fig. 4.**
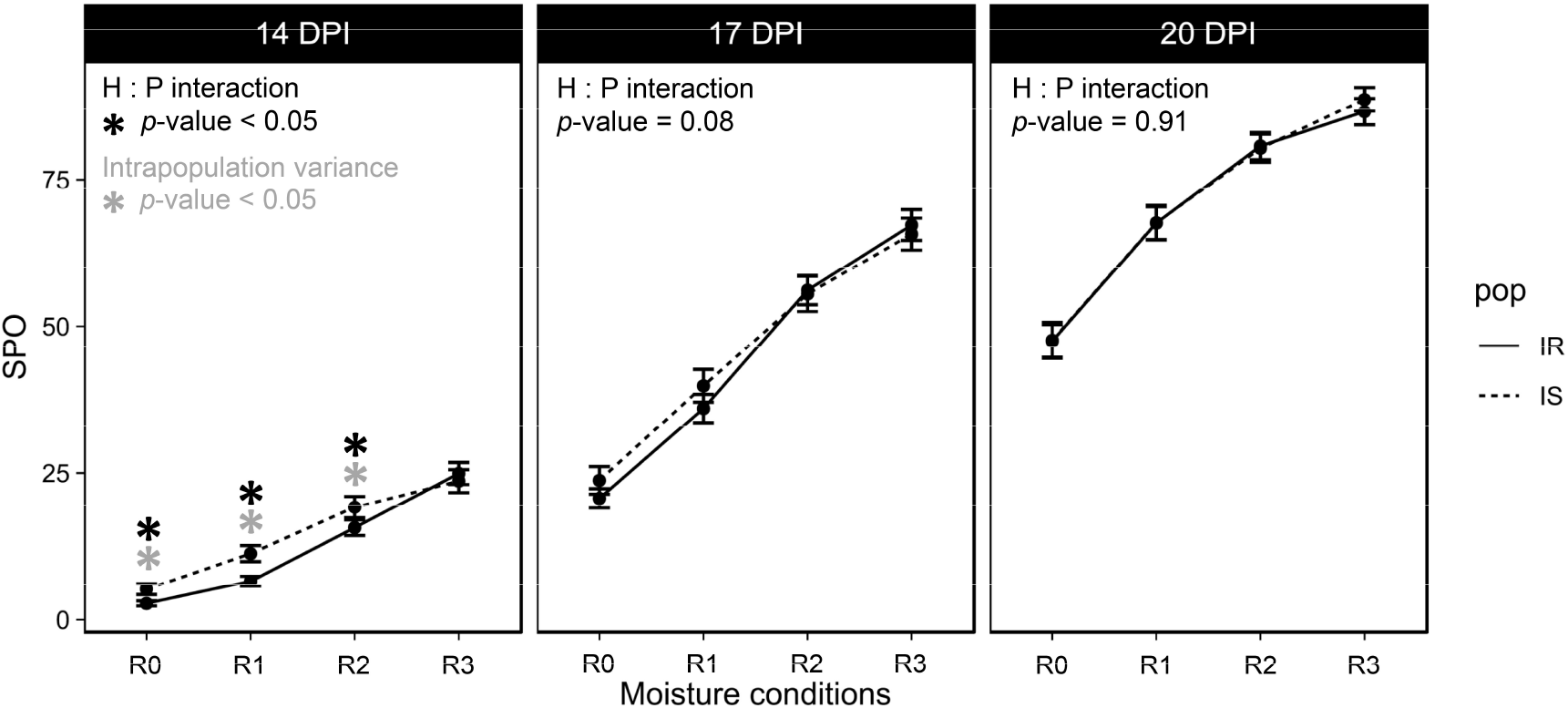
Dynamics of the effect of moisture regime on the percentage of sporulating area (SPO expressed as the mean ± SEM) in the early stages of *Zymoseptoria tritici* infection, by isolate origin - Ireland (IR; solid line) or Israel (IS; dashed line) - and the timing of disease assessment (14, 17, 20 dpi). Each data point corresponds to the mean of 24 isolates where each isolate is described by SPO assessments scores conducted on 6 wheat leaves taken from 2 pots placed in different growth chamber positions (opposite ends). Significant differences in the mean (*post-hoc* analysis of the GLM H×P interaction, *i.e.* the interaction of moisture regimes and differential population-level response to these conditions) and the intrapopulation variance (Levene’s test for homogeneity of variance) of SPO between IR population (*n* = 24 isolates) and IS population (*n* = 24 isolates) are highlighted by black and gray stars, respectively (*p*-value < 0.05). Moisture regimes: R0 = no-bagging treatment; R1 = one-day-bagging treatment; R2 = two-day-bagging treatment; R3 = three-day-bagging treatment.

### Significant effect of moisture during the early stages of infection

The temporal dynamics of the effect of moisture regimes justified the sequential fitting of GLMs at 14, 17, and 20 dpi (Table 1): at 14 dpi, the differences in sporulating area between R0-R1 and R2-R3 highlighted a dichotomy between unfavorable and favorable RH conditions. At 17 dpi, the mean differences between the four moisture regimes R0, R1, R2, and R3 were all significant (R0 ≠ R1 ≠ R2 ≠ R3). At 20 dpi, there were no longer any differences between R2 and R3, due to saturation (100%) being reached for sporulating area (see letters in Figure 3). From 0 to 3 dpi, the longer the duration of time at which infected wheat seedlings were exposed to a continuous RH 100%, the more rapid the development of lesions. Indeed, a slight difference in high moisture regime during the very early stages of *Z. tritici* infection, despite the relatively high RH value (80%) of the air saturation interruption period, had a strong impact on the dynamics of lesion development.

### Comparisons of tolerance to drier conditions are best performed at 14 dpi

Sporulating areas at 14 dpi were significantly larger for IS isolates on wheat plants subjected to the R0-R1-R2 regimes, whereas no significant difference was observed for R3 (Figure 4), highlighting the greater tolerance of drier conditions in the IS population. No difference was observed between the populations at 17 and 20 dpi (no statistically significant H×P interaction, although this interaction was close to significance at 17 dpi; GLM, *p*-value < 0.1), suggesting that the driest regimes delayed infection processes but did not prevent *Z. tritici* development. These results led us to perform further analyses at 14 dpi, through comparisons of ‘moisture reaction norms’ or ‘response curves’, to assess the differences between isolates within these two pathogen populations in more detail.

### Interindividual differences in moisture adaptation within the two populations at 14 dpi

The comparison of individual reaction norms (Figure S1), particularly for density distributions, and the ranges of values for the three parameters capturing the general characteristics of the response (intercept, slope, midpoint), highlighted high levels of interindividual variation in sensitivity to moisture regimes (Figure 5). Mean phenotypes did not differ significantly between the two populations (Wilcoxon rank sum test; differences in mean response assessed for the intercept: *p*-value = 0.33; slope: *p*-value = 0.39; midpoint: *p*-value = 0.92). For the population-level linear moisture reaction norm, a significant difference in sporulating area was observed between the IS and IR populations for R1 (Wilcoxon rank sum test, *p*-value = 0.03), but not for the other three moisture regimes (*p*-value > 0.05). This suggest that RH conditions may have been too restrictive or detrimental for the expression of biological differences and variance for the trait under the R0 regime, whereas they were not limiting for R2 and R3.

**Fig. 5.**
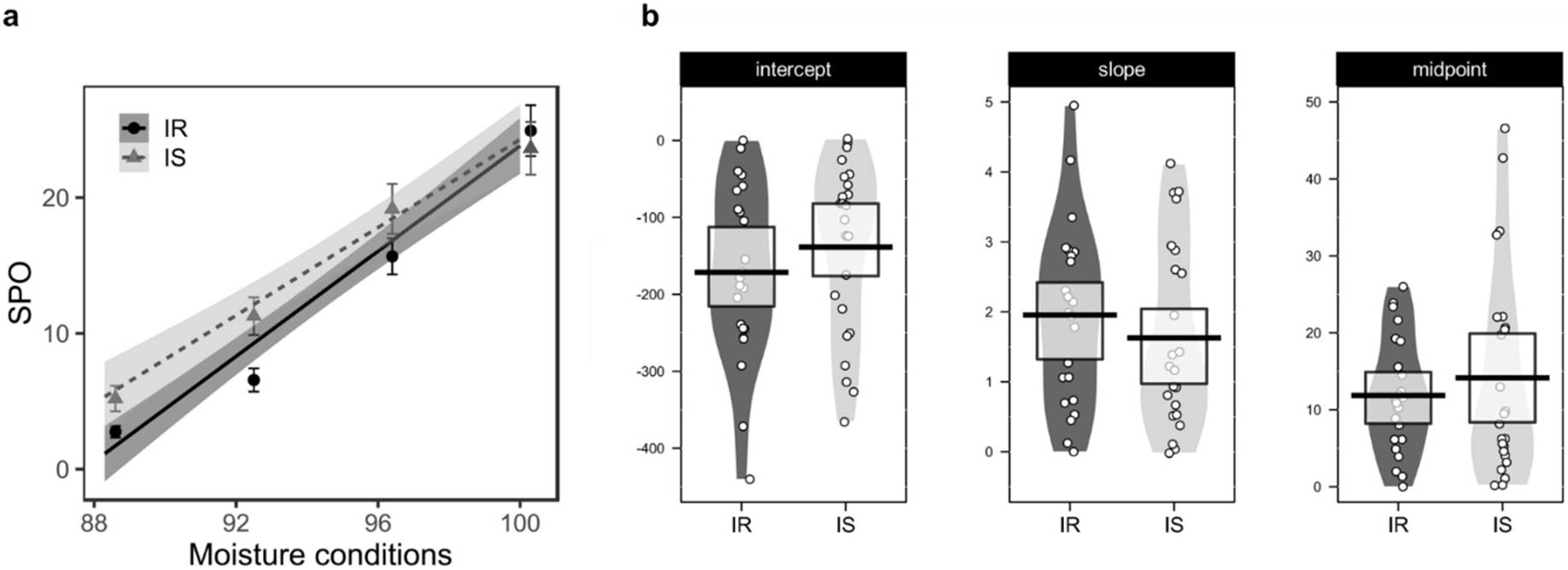
Variation of *Zymoseptoria tritici* response to moisture conditions at the population (IR: Irish population, IS: Israeli population) and individual levels, at 14 dpi. (a) Population-level moisture reaction norms (mean RH prevailing over the first three days after inoculation: 88.3% for R0, 92.2% for R1, 96.1% for R2, 100% for R3) for the IR (*n* = 24 isolates; black circles/solid line) and IS (*n* = 24 isolates; gray triangles/dashed line) populations. The corresponding linear regression lines SPO = a × RH + b (where a and b are the population average slope and intercept, respectively: a = 1.9 and b = −169.9 for IR; a = 1.6 and b = −137.6 for IS) fitted to the percentage of sporulating area (SPO, expressed as the mean ± SEM) are displayed with their 95% confidence intervals. (b) Interindividual variation of the three chosen descriptors of individual moisture reaction norms: intercept (*b*), slope (*a*) and midpoint (y_94.2%_). Individual values (open circles), means (horizontal black thick lines), distributions (smoothed density curves) and 95% Bayesian highest density intervals (central rectangular boxes enclosing the means) are shown for each descriptor for which interindividual variation is displayed by population (IR, IS; six biological replicates per isolate).

Neutral markers highlighted no differences in genetic structure between the IR and IS populations (Figure S2). P_ST_ values (computed at the critical c/h² ratio of 1) and their confidence intervals at c/h^2^ = 1 indicated a robust difference between P_ST_ and F_ST_ (Figure S3). Thus, any phenotypic difference suggests that interindividual differences in moisture adaptation within the two populations may conceal signatures of local adaptation.

### Classification of *Z. tritici* sensitivity to the four moisture regimes

Based on a PCA of phenotypic variation, we were able to classify the 48 *Z. tritici* isolates according to their sensitivity to the four moisture regimes (Figure 6). At 14 dpi, the R2 and R3 regimes did not discriminate between individual responses in terms of sporulating area. Cluster 4 consisted of isolates with a particularly large sporulating area (extreme phenotypes of is16, is19 and is34; Figure S1) that were less affected by the R0 and R1 regimes.

**Fig. 6.**
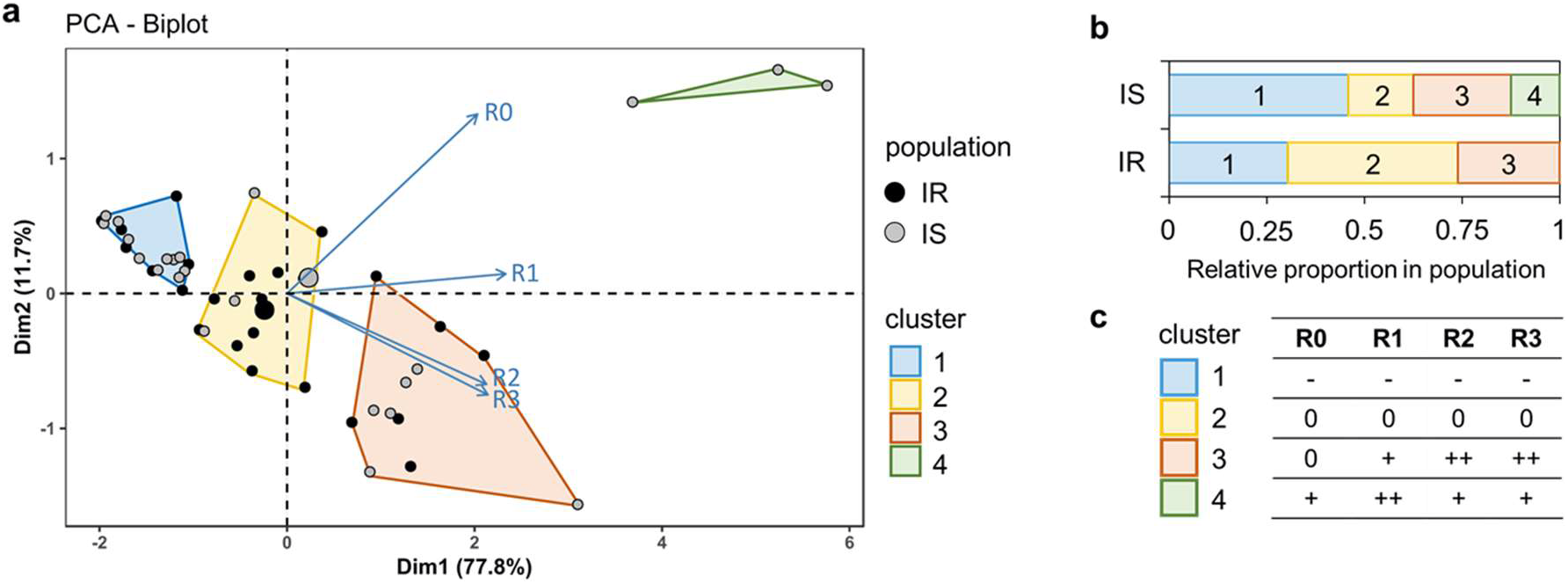
Classification of *Zymoseptoria tritici* sensitivity to the four moisture regimes at 14 dpi. (a) Principal component analysis (PCA) biplot showing Irish (IR, *n* = 24 isolates, black points) and Israeli (IS, *n* = 24 isolates, gray points) isolates plotted in two dimensions, using their projections onto the first two principal components to summarize their response (SPO, *i.e.* percentage of sporulating area) under the four moisture regimes. HCPC clusters of individual responses are shown as colored areas on the factorial plane. (b) Relative proportions of the four identified clusters in the IR and IS populations. (c) Reading grid for the clustering of isolates based on their responses to the four moisture regimes (six biological replicates per isolate): SPO significantly lower (−) or higher (+) than the dataset mean (0). Moisture regimes: R0 = no-bagging treatment; R1 = one-day-bagging treatment; R2 = two-day-bagging treatment; R3 = three-day-bagging treatment.

## Discussion

In this experimental study we highlighted the effect of moisture conditions during the early stages of *Z. tritici* infection: we established individual- and population-level moisture reaction norms, expressed as the area of *Z. tritici* sporulation on inoculated leaves for a RH averaged over the three days following inoculation. By this approach, we were able to quantify the critical effects of moisture conditions during the earliest stages of infection on the development of Septoria tritici blotch despite the relatively high RH value of the lowest moisture regime tested (R0). It also made it possible to refine the findings of several previous studies performed in our range of experimental conditions (Holmes, 1974; Shearer & Zadoks, 1972; Eyal, 1977; Hess *et al.*, 1987; Chungu *et al.*, 2001; Magboul *et al.*, 1992; Shaw, 1990; Shaw, 1991; Fones *et al.*, 2017). The effects of moisture conditions were more pronounced at 14 dpi than at 17 and 20 dpi. Based on these results, we analyzed the data acquired at 14 dpi in more detail, to determine moisture sensitivity at both the individual and population levels. These findings also raise the question as to why the effects of moisture conditions are less pronounced at 17 and 20 dpi, suggesting that the tolerance may simply be expressed as a ‘delay’ in the infection process, thus impacting the latent period. The higher tolerance of the Israeli population to early interruption of optimal moisture conditions disappeared after 20 dpi while the disease severity exceeded 75%, but such disease levels are rarely observed under field conditions. Thus, it cannot be excluded that what appears here to be a ‘delay’ is also an artefact related to the use of inoculum concentrations (10^5^ spores.mL^−1^) much higher than in droplets under field conditions, but nevertheless lower than in most of other experimental studies on *Z. tritici* (usually 10^6^ or 10^7^ spores.mL^−1^). The use of lower inoculum concentrations in further experiments to avoid saturation effects would lead to the appearance of small non-coalescent lesions and could thus improve the interpretation of the results. Fones *et al.* (2017) previously showed that increasing leaf moisture is associated with increasing disease severity (number of pycnidia per leaf), because it decreases the time required for *Z. tritici* to penetrate into leaf tissues. This finding is consistent with our results, suggesting that moisture conditions have a greater impact on the rate at which symptoms appear than on their final intensity. We may hypothesize that suboptimal moisture regimes have slowed epiphytic hyphal growth and penetration into the leaf tissues via the stomata, but that they did not ultimately reduce the germination rate or infection efficiency, at least for the RH values which were tested here. Epiphytic development was probably slower during the daytime periods, when wheat leaves were not bagged, resulting in a ‘delay’ in the infection process rather than an ‘irreversible’ ending of this process. This hypothesis should be tested with direct experimental approaches based on cytological observations or indirect approaches measuring infection efficiency, defined as the proportion of pathogen spores able to infect susceptible plant tissues once they have landed on them.

As previously reported, the impact of RH cannot be tested alone without considering the effect of liquid water, both variables being linked by dew point. In all conditions, excepted 80% RH at 22°C, the dew point was theoretically reached (despite liquid water was observed only on the bag surface and the walls of the chamber) but it was not possible to verify it. This uncertainty could have been avoided: (i) by working deliberately at 99% RH rather than at saturation; (ii) by modulating the temperature in each moisture condition, but this would have required more sophisticated technical facilities; (iii) by using a ‘cold sink’ above the plants (e.g. produced by freezing a suitable substance at −15°C or −20°C) that would lead to the leaves being marginally cooler than the surrounding air reproducing outdoor clear night sky conditions. Moreover, covering plants may have resulted in significant changes in other climatic factors within the bag, such as leaf temperature: we definitely should be aware of the strong complex relationship between moisture – RH and liquid water on leaf – and temperature. Continuing to study the impact of one of these factors on the development of plant pathogens involves looking at the other one. The current study, the first one on pathogen adaptation to moisture, highlights how this question is worth pursuing. Further *in planta* experiments should consider the interactions between climatic factors and ensure their fine control in experimental facilities using for instance dedicated moisture sensors and thermocouples, positioned in the air but also in direct contact with the leaf.

Even under controlled conditions, growth chambers are subject to spatiotemporal fluctuations in within-chamber conditions, particularly as concerns the distribution of light intensity (Potvin *et al*., 1990). In general, such spatial patterns of heterogeneity constitute a difficult problem to solve when working *in planta* which justifies to use appropriate experimental designs taking them into account (see the block effect; Table 1). Differences in inoculation conditions might also explain the differences in symptom expression between series. One possible improvement to the experimental method would be to screen larger numbers of isolates in a more standardized way (making it easier to separate environmental and physiological responses), for instance by controlling and testing moisture conditions *in vitro* using the experimental devices proposed by Li *et al.* (2014; different RH conditions formed within humidity chambers obtained with different glycerol solution concentrations, even though saturated saline solutions might be preferable due to the more stable relative humidity they induce) or Xu *et al.* (2016; discrete RH gradient formed in spatially separated wells of a multi-well plate). This would have the advantage of countering the aforementioned technical difficulties and the effect of the host resistance × environmental response interaction on the expression of pathogenicity in isolates, although, in our experiment, the two *Z. tritici* populations appeared similarly aggressive on the cultivar ‘Taichung 29’. *In vitro* approaches may be particularly relevant for *Z. tritici*, as they have been shown to be a reliable *proxy* for the assessment of differences in thermal sensitivity and allowed to test many more strains and populations (Boixel *et al*., 2019a). However, as an *in vitro* response corresponds to an ‘abnormal’ physiological state of the fungus in a simplified environment, extrapolation to field conditions can become more of an epistemological than a technical issue.

To our knowledge, this experimental study provides the first evidence of moisture adaptation in a fungal plant pathogen. The results in Figure 5 highlight strong individual variations in the phenotypic plasticity of *Z. tritici* with respect to sensitivity to high moisture regimes. Notwithstanding the relatively limited sample size (48 isolates) and number of populations (*n* = 2) investigated here, this study revealed, for the first time in a fungal plant pathogen, the existence of individual variation in responses to RH conditions. Moreover, the Israeli population was shown to be more tolerant to early interruption of optimal moisture conditions. Together with the absence of genotypic differentiation for neutral microsatellite loci between the two populations, this result reveals signature of adaptation to moisture conditions which is consistent with the climatic conditions of the areas from which the two populations were collected.

The interaction between RH and isolate effects was significant whenever disease was assessed (see RH conditions × isolate interaction (H×I(P)) in Table 1), highlighting the importance of taking the interindividual variation in response to moisture conditions into account. Differences in moisture sensitivity were analyzed at both the population and individual levels. The moisture performance curves highlighted sensitivity variations between the two populations. The differences in the mean values of the parameters characterizing this sensitivity were not significant. However, the interindividual analysis revealed significant differences in moisture sensitivity between isolates, suggesting that it would be possible to quantify such differences more accurately in future experiments. This would require the study of larger numbers of individuals and more stringent moisture conditions. For instance, lower moisture regimes could be tested to provide an accurate analysis of moisture adaptation to lower humidity levels which are similar to the annual air relative humidity conditions experienced by the Israeli population (69.7%).

The Israeli and Irish *Z. tritici* populations studied here are part of a Euro-Mediterranean set of eight populations included in a broader study on thermal adaptation (Boixel *et al.*, 2019b). General ecological concepts, knowledge and methods have been developed to a greater extent for thermal biology than for moisture biology. The estimation of interactions between moisture and temperature adaptation would also be relevant at both population and individual level. For this purpose, *Z. tritici* is an interesting case, both because a physical relationship between the two microclimatic variables, temperature and moisture, has been established (Shaw & Royle, 1993; Pietravalle *et al.*, 2003), and because the development of Septoria tritici blotch is known to be strongly influenced by both variables. The interactions between temperature and moisture effects could be studied from a mechanistic angle (for instance, testing pleiotropy *vs*. co-selection hypotheses in the genetic determinism of such a dual adaptation) but also from an epidemiological angle (predicting the consequences of climate changes affecting plant disease development in a more holistic way). The comparison of differences between temperature and moisture responses in different pathogen populations, starting with the Israeli and Irish populations, is particularly interesting because moisture adaptation is already interconnected with thermal adaptation in some models (e.g. Shinozaki & Yamaguchi-Shinozaki, 2000).

This study did not aim to determine the impact of mean RH value (constant moisture regime) nor the RH threshold below which *Z. tritici* isolates cannot infect wheat, but we can unambiguously conclude that “a moisture regime as close as possible to air saturation is best for *Z. tritici*”. The minimal RH threshold for *Z. tritici* remains unknown. The challenge of answering this question is however quite limited since the moisture conditions to which a foliar pathogen is actually exposed in the field fluctuate, justifying the relevance of focusing on responses to fluctuant rather than constant regimes. For instance, Shaw (1991) showed that breaks at 50% relative humidity had large effects but nevertheless enabled infection to occur. We can assume that there is a threshold moisture level (a lethal condition definitively blocking the infectious process), like that for temperature, but previous studies in *Z. tritici* and general knowledge about the epidemiology of fungal disease suggest that a minimum threshold, rather than a maximum, must be taken into account (as RH, by definition, cannot exceed 100%), together with a time period during which RH remains below this minimum threshold. Conversely, for temperature, the lethal threshold that should be considered is a maximum value, because *Z. tritici* can survive the thermal conditions of winter and freezing at −80°C in laboratory conditions.

Comparing the effects of the duration of uninterrupted periods of exposure to optimal moisture conditions as it was done here (see also Shaw, 1991; Magboul *et al.*, 1992; Chungu *et al.*, 2001) rather than effects of mean RH values (Fones *et al*., 2017) makes the comparison of results from different experimental studies more complex. Therefore, one might be tempted to compare directly constant moisture conditions (e.g. optimal 100% RH *vs.* limiting 50% RH). However, this poorly represents what is happening in the field where fluctuating rather than constant moisture conditions prevail. We must keep in mind that average RH (e.g. 78-92% of RH in Carlow, Ireland; 67-73% of RH in Kiryat-Tivon, Israel; Figure 1) is a simplified description: even in the driest areas, the level of moisture in a wheat canopy can be very high during or just after a rain period regardless of the smoothing average monthly air RH recorded at a site, and a low daytime RH is not inconsistent with dew under a clear sky and with a well-watered crop. Nevertheless, the moisture regimes tested in this study are a step closer to better understand the response to actual field conditions for which canopy RH is maximal during the night and lower during the day. More generally, the question of how best to describe RH conditions remains unresolved: averaging, intermittent favorable/unfavorable conditions, etc. How can the effect of the putative minimum moisture threshold (still unknown) be linked to the effect of the duration of dry periods? To which microclimatic variables are the individuals and populations really (mal)adapted? These questions and the spatio-temporal resolution of measurements should be addressed. For instance, it has been experimentally demonstrated that *Z. tritici* responds to ‘leaf temperature’ (mirroring with ‘body temperature’ for animals) rather than ‘air temperature’ (Bernard *et al.*, 2013), and that the time step for measurement is important. These conclusions could also be likely extended to moisture.

Additional studies on adaptation to very suboptimal moisture regimes (e.g. 50-60% RH) rather than regimes close to the optimum (e.g. 90-100% RH) should be considered: (i) for methodological reasons, because this would probably maximize the expression of adaptation, making it easier to characterize experimentally, with populations contrasting less than the Israeli and Irish populations studied here; (ii) for epidemiological reasons, because it would provide knowledge to improve predictions of the adaptation of *Z. tritici* populations on bread and durum wheat in dryland farming areas (e.g. in Middle East and North Africa) in response to climate change. Such aims are ambitious and the studies will need to take into account the difficulties involved in experimental studies investigating the effects of moisture.

The exploitation of relevant analyses based on biological data requires attention to the issue of ‘moisture stress tolerance’ and an extension of reflections to the ecology of communities (Sheik *et al.*, 2011). This study provides preliminary insight into such moisture stress tolerance and the diversity of its responses across individuals and populations when considering the effects of climate change on plant disease epidemics. A knowledge of the microclimatic requirements for fungal pathogen development is essential for prospective studies of the impact of climate change with a solid experimental basis. To date, such studies have mostly focused on temperature, but climate change in the global context will also include large changes in moisture conditions with pronounced domino effects.

## Data Availability Statement

The data that support the findings of this study are openly available in the INRAE Dataverse online data repository (https://data.inrae.fr/) at https://doi.org/10.15454/FK7WHW.

## Author Contributions

FS conceived the study and was in charge of overall direction and planning, with substantial input from ALB. SG performed the experiments according to a protocol developed jointly by FS and TM. ALB and FS analyzed the data, prepared the figures and wrote the manuscript. TM provided critical feedback. All authors approved the final version of the manuscript.

## Acknowledgements

We are grateful to Dr. Stephen Kildea (Teagasc, Agriculture and Food Development Authority) and Dr. Hanan Sela (Institute for Cereal Crops Improvement, Tel Aviv University) for their help in sampling the Irish and Israeli *Z. tritici* populations, to Martin Willigsecker (INRAE BIOGER) for technical support throughout the experiment, and to Dr. Julie Sappa for her editorial advice in our English usage.

## Supplementary Material

Supplementary material is available in the online version of this article.

---

**Fig. S1.**
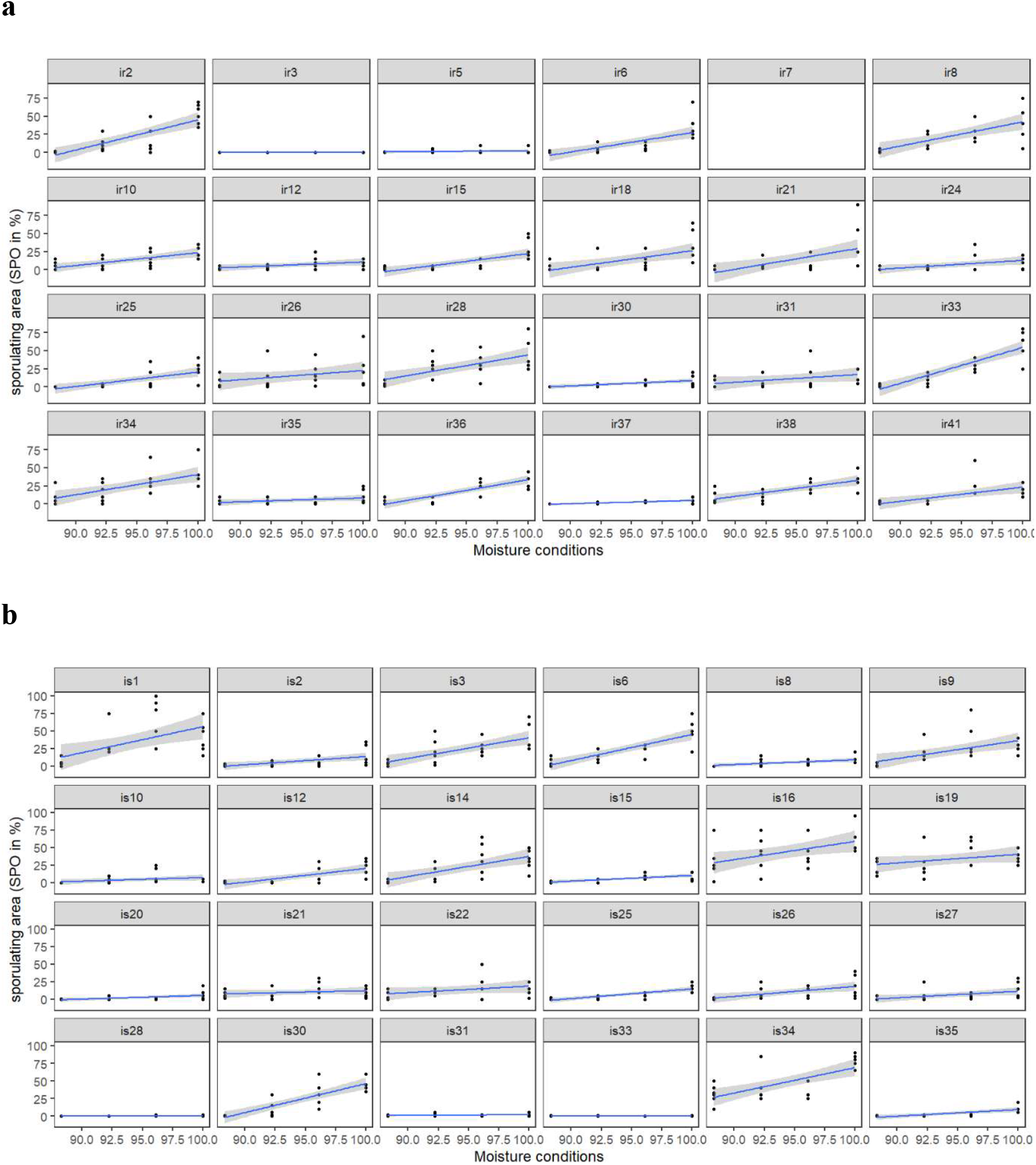
Individual moisture reaction norms at 14 dpi for (a) the 24 Irish (IR) and the (b) 24 Israeli (IS) *Zymoseptoria tritici* isolates. Sporulating areas (SPO) are expressed as a percentage of the inoculated leaf area for each isolate (except ir7, due to unsuccessful inoculation), in response to four moisture conditions characterized by the mean RH prevailing during the three-day period immediately after inoculation (see Figure 2). Linear regression lines (in blue) and their 95% confidence intervals were fitted independently to experimental observations (dark points corresponding to six seedling wheat leaves per isolate × moisture conditions, *i.e.* six biological replicates).

**Fig. S2.**
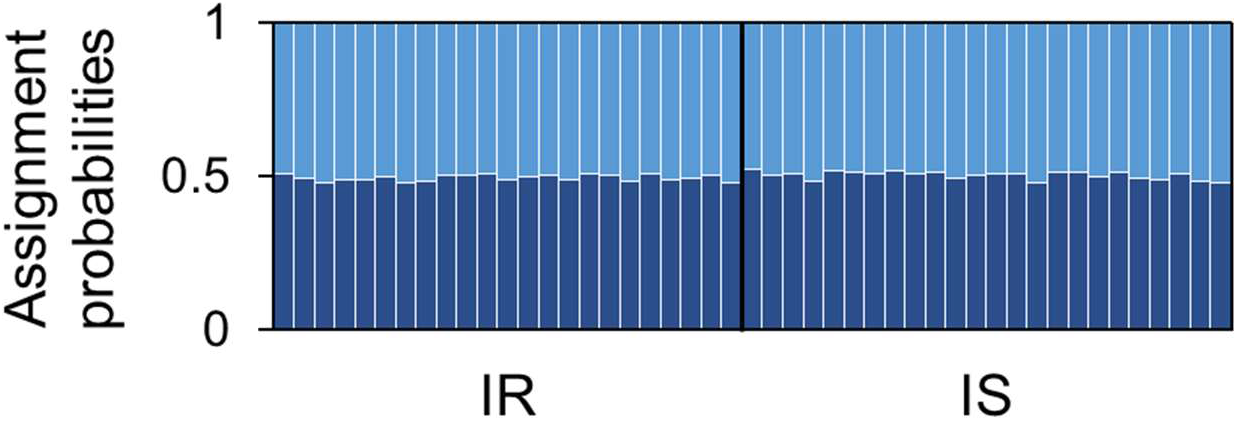
Genetic subdivision of the 48 *Zymoseptoria tritici* isolates composing the Irish (IR) and Israeli (IS) populations. Bar plot displaying the Bayesian genetic clustering of IR and IS populations according to 12 SSR markers. The STRUCTURE algorithm was applied under the admixture and correlated allele frequencies model (500,000 iterations of the Markov chain followed by a run phase of 1,000,000 iterations with 10 independent replicates for each tested number of clusters). Each isolate is represented by a single vertical bar broken into two color segments, the lengths of which are proportional to the probability of to the isolate being assigned to the inferred genetic clusters (*K* = 2). No population structure was detected because each individual was affected to the three genetic clusters with similar probabilities.

**Fig. S3.**
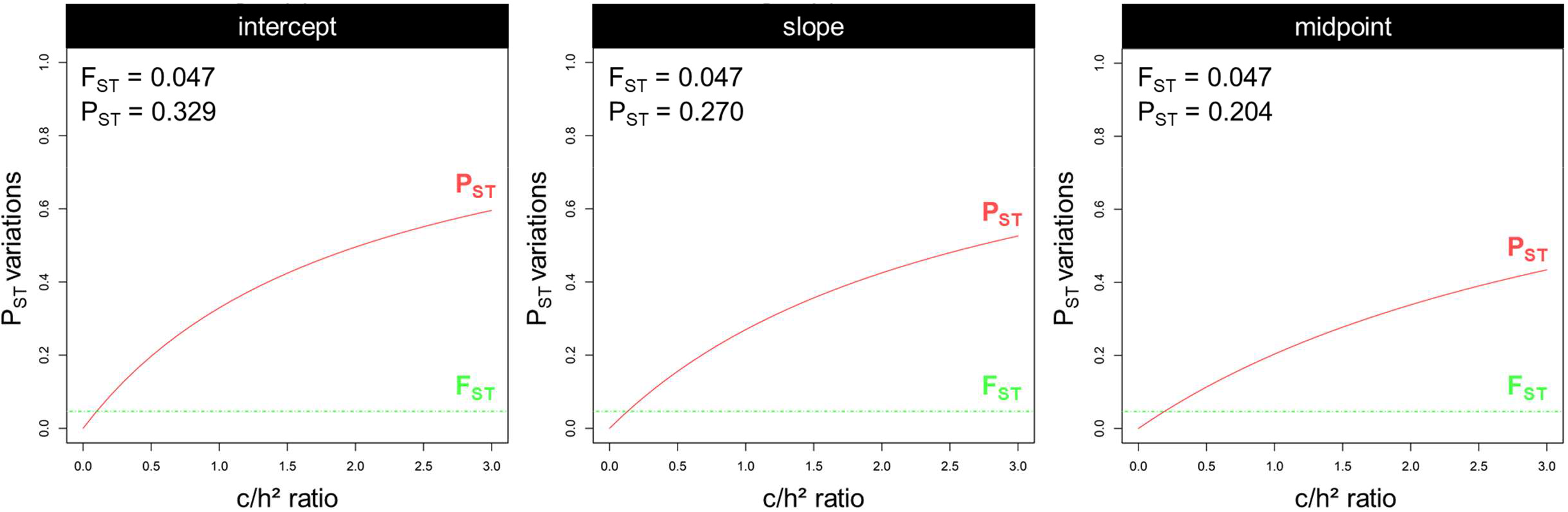
Phenotypic divergence in moisture reaction norms between the IR and IS populations (*n* = 48 isolates). Sensitivity analyses exploring the robustness of P_ST_–F_ST_ comparisons over variations in the c/h^2^ ratio, which determines the accuracy of the approximation of Q_ST_ by P_ST_. Phenotypic divergence in individual moisture reaction norms between populations (P_ST_) was investigated for the intercept, slope and midpoint parameters. On each plot, estimates of P_ST_ (solid line in red) and F_ST_ (dashed line in green) are shown, to reveal the contrasting patterns of phenotypic responses to moisture conditions and neutral genetic divergence. The P_ST_ values displayed (computed at the critical c/h² ratio of 1) and their confidence intervals at c/h^2^ = 1 indicate the occurrence of a robust difference between P_ST_ and F_ST_, suggesting the existence of signatures of local adaptation.

## References

Bernard, F., Sache, I., Suffert, F., Chelle, M. (2013). The development of a foliar fungal pathogen does react to leaf temperature! New Phytologist 198, 232–240.

Boixel, A.-L., Chelle, M., Suffert, F. (2019b). Patterns of thermal adaptation in a worldwide plant pathogen: local diversity and plasticity reveal two-tier dynamics. bioRxiv https://doi.org/10.1101/867572.

Boixel, A.-L., Delestre, G., Legeay, J., Chelle, M., Suffert, F. (2019a). Phenotyping thermal responses of yeasts and yeast-like microorganisms at the individual and population levels: proof-of-concept, development and application of an experimental framework to a plant pathogen. Microbial Ecology 78, 42–56.

Campbell, G.S., Norman, J.M. (1998). An introduction to environmental biophysics, 2nd ed. Springer-Verlag, New York, NY, USA.

Chakraborty, S., Tiedemann, A.V., Teng, P.S. (2000). Climate change: potential impact on plant diseases. Environmental Pollution 108, 317–326.

Chungu, C., Gilbert, J., Townley-Smith, F. (2001). *Septoria tritici* blotch development as affected by temperature, duration of leaf wetness, inoculum concentration, and host. Plant Disease 85, 430–435.

Dawson, T.E., Goldsmith, G.R. (2018). The value of wet leaves. New Phytologist 219, 1156–1169.

Duncan, K.E., Howard, R.J. (2000). Cytological analysis of wheat infection by the leaf blotch pathogen *Mycosphaerella graminicola*. Mycological Research 104, 1074–1082.

Eyal, Z., Brown, J.F., Krupinsky, J.M., Scharen, A.L. (1977). The effect of postinoculation periods of leaf wetness on the response of wheat cultivars to infection by *Septoria nodorum*. Phytopathology 67, 874–878.

Fones, H.N., Eyles, C.J., Kay, W., Cowper, J., Gurr, S.J. (2017). A role for random, humidity-dependent epiphytic growth prior to invasion of wheat by *Zymoseptoria tritici*. Fungal Genetics and Biology 106, 51–60.

Garrett, K.A., Dendy, S.P., Frank, E.E., Rouse, M.N., Travers, S.E. (2006). Climate change effects on plant disease: genomes to ecosystems. Annual Review of Phytopathology 44, 489–509.

Hess, D.E., Shaner, G. (1987). Effect of moisture and temperature on development of septoria tritici blotch in wheat. Phytopathology 77, 215–219.

Holmes, S.J.I., Colhoun, J. (1974). Infection of wheat by *Septoria nodorum* and *S. tritici* in relation to plant age, air temperature and relative humidity. Transactions of the British Mycological Society 63, 329–338.

Huber, L., Gillespie, T.J. (1992). Modeling leaf wetness in relation to plant‐disease epidemiology. Annual Review of Phytopathology 30, 553–577.

Huntingford, C., Jones, R.G., Prudhomme, C., Lamb, R., Gash, J.H., & Jones, D.A. (2003). Regional climate‐model predictions of extreme rainfall for a changing climate. Quarterly Journal of the Royal Meteorological Society 129, 1607–1621.

Jhorara, O.P., Butlerb, D.R., Mathaudaa, S.S. (1998). Effects of leaf wetness duration, relative humidity, light and dark on infection and sporulation by *Didymella rabiei* on chickpea. Plant Pathology 47, 586–594.

de Jong, G. (1995) Phenotypic plasticity as a product of selection in a variable environment. American Naturalist 145, 493–512.

Li, Y., Uddin, W., Kaminski, J.E. (2014). Effects of relative humidity on infection, colonization and conidiation of *Magnaporthe orzyae* on perennial ryegrass. Plant Pathology 63, 590–597.

Magboul, A.M, Geng, S., Gilchrist, D.G., Jackson, L.F. (1992). Environmental influence on the infection of wheat by *Mycosphaerella graminicola*. Phytopathology 82, 1407–1413.

Makhdoomi, A., Mehrabi, R., Khodarahmi, M., Abrinbana, M. (2015). Efficacy of wheat genotypes and Stb resistance genes against Iranian isolates of *Zymoseptoria tritici*. Journal of General Plant Pathology 81, 5–14.

Merilä, J., Crnokrak, P. (2001). Comparison of genetic differentiation at marker loci and quantitative traits. Journal of Evolutionary Biology 14, 892–903.

Pachinburavan, A. (1981). Pycnidiospore germination, penetration, and pycnidial formation of *Septoria tritici* Rob. ex Desm. PhD thesis. Washington State University, Washington, USA.

Pietravalle, S., Shaw, M.W., Parker, S.R., Van Den Bosch, F. (2003). Modeling of relationships between weather and *Septoria tritici* epidemics on winter wheat: a critical approach. Phytopathology 93, 1329–1339.

Potvin, C., Lechowicz, M.J., Bell, G., Schoen, D. (1990). Spatial, temporal, and species-specific patterns of heterogeneity in growth chamber experiments. Canadian Journal of Botany 68, 499–504.

Rowlandson, T., Gleason, M., Sentelhas, P., Gillespie, T., Thomas, C., Hornbuckle, B. (2015). Reconsidering leaf wetness duration determination for plant disease management. Plant Disease 99, 310–319.

Sentelhas, P.C., Gillespie, T.J., Gleason, M.L., Monteiro, J.E.B., Helland, S.T. (2004). Operational exposure of leaf wetness sensors. Agricultural and Forest Meteorology 126, 59–72.

Shaw, M.W. (1990). Effects of temperature, leaf wetness and cultivar on the latent period of *Mycosphaerella graminicola* on winter wheat. Plant Pathology 39, 255–268.

Shaw, M.W. (1991). Interacting effects of interrupted humid periods and light on infection of wheat leaves by *Mycosphaerella graminicola* (*Septoria tritici*). Plant Pathology 40, 595–607.

Shaw, M.W., Royle, D.J. (1993). Factors determining the severity of epidemics of *Mycosphaerella graminicola* (*Septoria tritici*) on winter wheat in the UK. Plant Pathology 42, 882–899.

Shearer, B.L., Zadoks, J.C. (1972). The latent period of *Septoria nodorum* in wheat. 1. The effect of temperature and moisture treatments under controlled conditions. Netherlands Journal of Plant Pathology 78, 231–241.

Sheik, C.S., Beasley, W.H., Elshahed, M.S., Zhou, X., Luo, Y., Krumholz, L.R. (2011). Effect of warming and drought on grassland microbial communities. The ISME Journal 5, 1692–1700.

Shinozaki, K., Yamaguchi-Shinozaki, K. (2000). Molecular responses to dehydration and low temperature: differences and cross-talk between two stress signaling pathways. Current Opinion in Plant Biology 3, 217–223.

Suffert, F., Ravigné, V., Sache, I. (2015). Seasonal changes drive short-term selection for fitness traits in the wheat pathogen *Zymoseptoria tritici*. Applied and Environmental Microbiology 81, 6367–6379.

Suffert, F., Thompson, R. (2018). Some reasons why the latent period should not always be considered constant over the course of a plant disease epidemic. Plant Pathology 67, 1831–1840.

Weiss, A. (1990). Leaf wetness: Measurements and models. Remote Sensing Reviews 5, 215–224.

West, J.S, Townsend, J.A., Stevens, M., Fitt, B.D.L. (2012). Comparative biology of different plant pathogens to estimate effects of climate change on crop diseases in Europe. European Journal of Plant Pathology 133, 315–331.

Xu, Z., Jiang, H., Sahu, B.B., Kambakam, S., Singh, P., Wang, X., Wang, G., Bhattacharyya, M.K., Dong, L. (2016). Humidity assay for studying plant-pathogen interactions in miniature controlled discrete humidity environments with good throughput. Biomicrofluidics 10, 034108.

